# Live-cell microscopy reveals that human T cells primarily respond chemokinetically within a CCL19 gradient that induces chemotaxis in dendritic cells

**DOI:** 10.1101/2020.11.08.373548

**Authors:** Evert J. Loef, Hilary M. Sheppard, Nigel P. Birch, P. Rod Dunbar

## Abstract

The ability to study migratory behavior of immune cells is crucial to understanding the dynamic control of the immune system. Migration induced by chemokines is often assumed to be directional (chemotaxis), yet commonly used endpoint migration assays are confounded by detecting increased cell migration that lacks directionality (chemokinesis).

To distinguish between chemotaxis and chemokinesis we used the classic “under-agarose assay” in combination with video-microscopy to monitor migration of CCR7+ human monocyte-derived dendritic cells and T cells in response to a concentration gradient of CCL19. The formation of the gradients was visualized with a fluorescent marker and lasted several hours.

Monocyte-derived dendritic cells migrated chemotactically towards the CCL19 gradient. In contrast, T cells exhibited a biased random walk that was primarily driven by increased exploratory chemokinesis towards CCL19. This dominance of chemokinesis over chemotaxis in T cells is consistent with CCR7 ligation optimizing T cell scanning of antigen-presenting cells in lymphoid tissues.

## 1 Introduction

Cell migration is a crucial process in a myriad of physiological functions (Comerford et al. 2013). Homing of both T cells and dendritic cells to lymph nodes is mainly dependent on activation of the chemokine receptor CCR7 (Comerford et al. 2013). Co-localization of these cells within the T cell zones present in lymph nodes allows T cells to scan DCs for their cognate antigen, ultimately enabling activation and expansion of antigen-specific T cells (Förster, Davalos-Misslitz, and Rot 2008). The CCR7 ligand CCL19 is considered to be strongly chemotactic for both T cells and dendritic cells (DCs), potentially driving their co-localization. However, intra-vital microscopy in mice has revealed that migration of T cells *within* lymph nodes does not have strong features of directional chemotaxis (Worbs et al. 2007). It has been suggested that the random directionality observed *in vivo* doesn’t exclude the possibility that there is an underlying directional bias (Bogle and Dunbar 2008). Understanding how CCL19 might act through the same receptor to generate different types of migratory behavior in T cells and DCs is central to understanding the dynamic control of T cell responses.

Data to support the concept that CCL19 drives chemotaxis for both T cells and DCs are often drawn from “transwell” assays that are based on the original Boyden assay (Pujic et al. 2009). In these assays, the cells and the chemotactic agent are separated by a membrane with pores large enough to permit cell migration; the number of cells that move from the cell chamber to the second chamber are counted. Given that the pores in the barriers are large enough to permit cell transit, it is likely that chemotactic agents applied to one chamber will rapidly equilibrate in the other chamber. Some researchers have modified these assays by coating the porous barriers with extracellular matrix, fibrin or collagen gels, and in some cases, monolayers of endothelial or epithelial cells. Under these conditions, concentration gradients may be maintained for periods long enough to assess cell migration. However, these transwell assays are typically used as endpoint assays so crucial migratory information is not measurable, such as migration speed and track straightness toward the chemokine over time. The use of endpoint assays also introduces a confounding error in terms of measuring directional chemotaxis: agents that simply increase the speed of migration of cells, with no or minimal directional component, will increase the number of cells detected in the second chamber, effectively reading out chemokinetic effects or biased random walks as chemotaxis.

The study presented here aimed to use a simple real-time migration assay that would allow for the detailed analysis of migration of human mDCs and T cells in response to chemokines.

We used the “under-agarose” assay that was initially developed in 1975 by Simmons *et al*. (Nelson, Quie, and Simmons 1975). This assay allows the researcher to set up two (or more) competing chemoattractant signals whereby chemoattractants diffuse slowly through gels rather than rapidly equilibrating in solution. The presence of a gel also allows for the study of cell movement in a confined plane, allowing for an integrin-independent amoeboid type of migration that mimics the primary kind of locomotion of DCs and T cells in 3D matrices, which is suggested to be better suited to rapidly follow chemotactic gradients. (Krummel, Friedman, and Jacobelli 2014; Lämmermann et al. 2008; Friedl et al. 1998). This assay, and other similar assays have been widely used to study the migration of cells (Heit and Kubes 2003; Vargas et al. 2016; Sixt and Lämmermann 2011; Visweshwaran and Maritzen 2019). In this study, we used agar rather than agarose as this increased the number of migrating cells. The use of live-cell microscopy enabled the visualization of the migration of human monocyte-derived dendritic cells (mDCs) and human T cells in real-time. Because CCL19, unlike CCL21, is a soluble chemokine (Barmore et al. 2016), Fluorescent dextrans of a similar size (10 kDa) were used to demonstrate that a concentration gradient was generated that lasted for several hours. This provided sufficient time to allow for definitive tracking of cell migration paths over hundreds of microns in the presence of a CCL19 gradient or a uniform CCL19 concentration.

This method showed that human mDCs exhibit true chemotaxis toward a gradient of CCL19. Human polyclonal T cells, however, respond to a CCL19 gradient with a biased random walk, showing directional bias, but mostly chemokinetic, and showing similarities to the response to a uniform CCL19 concentration. The strong chemokinetic response in T cells is consistent with efficient strategies to scan antigen-presenting cells for cognate antigen within lymphoid tissue.

## 2 Materials and methods

### Cell culture

All cytokines were purchased from Peprotech (Rocky Hill, NJ, USA).Human blood was obtained from healthy volunteers after informed consent and with approval by the University of Auckland Human Participant Ethics Committee (Ethics Approval 010558). Peripheral blood mononuclear cells were prepared using Lymphoprep (Axis-Shield, Dundee, Scotland) density gradient centrifugation. mDCs were differentiated from CD14^+^ monocytes based on a previously reported method (Lehner et al. 2005). In short, CD14^+^ cells were isolated using the MACS human CD14^+^ isolation kit (Miltenyi Biotec, Bergisch Gladbach, Germany) according to the manufacturer’s protocol. One to two *10^6^ CD14^+^ cells were plated in a 24 well plate with AIM-V medium (Life Technologies, Carlsbad, California, USA) supplemented with 1x GlutaMAX (Life Technologies) and 200 ng mL^−1^ IL-4 and 100 ng mL^−1^ GM-CMSF. Half of the medium was replaced at day 2 or 3. On day five, non-adherent and mildly-adherent cells were resuspended and transferred to a 15 mL conical tube and centrifuged at 350 g for 5 minutes. The pellet was resuspended in 1 mL fresh AIM-V containing 100 ng mL^−1^ GM-CSF, 10 ng mL^−1^ IL-1β, 100 ng mL^−1^ IL-6, 250 ng mL^−1^ TNF-a, and 1 μg mL^−1^ PGE2 to mature the cells for a further 48 hours. Expanded T cells, in this manuscript only referred to as T cells were cultured in RPMI-1640 medium containing 5% human serum (One Lambda, Los Angles, California, USA), 100 U mL^−1^ penicillin (Life Technologies), 100 μg mL^−1^ streptomycin (Life Technologies), and 2 mM GlutaMAX-1 (Life Technologies), supplemented with 5 ng mL^−1^ IL-7 (referred to as RS5-IL7) unless stated otherwise. T cells were polyclonally expanded from freshly isolated PBMCs using Dynabeads human T-activator CD3/CD28 beads (Life Technologies) as previously described (Loef et al. 2019; N. Lorenz et al. 2015; Natalie Lorenz et al. 2016). In brief, 1.10^6^ PBMCs were activated with Dynabeads at a bead:cell ratio of 1:1 for 3 days in RS5-IL7, supplemented with 10 ng mL^−1^ IL-12 and 10 ng mL^−1^ IL-21. Following magnetic removal of the beads, the cells were cultured for a further 4 days using the same medium, followed by 7 days in RS5-IL7 supplemented with IL-21. Cells were examined daily, and cultures were split once cells were confluent, or the medium showed signs of acidification (usually every 2-3 days). Cells were rested for a further 7-10 days in RS5-IL7 prior to use or cryopreservation. Cryopreserved T cells were allowed to recover for at least 24 h in RS5-IL7 (20 ng.mL^−1^) before use. Post recovery, every T cell expansion was tested for expression of CCR7 by flow cytometry, and only T cells expansions that showed a more than 50% CCR7^+^ population were used in experiments.

### Agar set up

To make 0.5% agar gels, 2 mL 2x RPMI (made from Powder) (Sigma St. Louis, Missouri, USA) was mixed with 200 μl human serum (One Lambda, Los Angles, California, USA) and 800 μl ultrapure H2O. This was prewarmed to 37°C in a water bath. 2% agar was dissolved in ultrapure H2O by bringing it to a boil in the microwave and mixing it on high speed on a vortex mixer for 20 seconds. This process was repeated four times. One mL of the agar solution was added to the prewarmed mixture to make a 0.5% agar medium solution. Of the solution, 800 μl was added to each well of a 4 well 1.5 polymer tissue culture treated chambered coverslip (Ibidi, Martinsried, Germany) that was precoated with 20% human serum in RPMI for 30 min at 37°C. To generate a uniform concentration of CCL19 (PeproTech), CCL19 was added to a final concentration of 100 ng mL^−1^ before letting the agar solidify. The agar was left to set for 1 hour. Next, a three-pronged bespoke autopsy punch was used to create a line of three wells, each of a three mm diameter and 2 mm apart in the agar (Supplementary Figure 1).

### Microscopy

The cells were stained with Cytotrack green or red (Bio-Rad, Hercules, California, USA). The dye was diluted 1:500 in PBS. The cells were resuspended in the PBS dye solution at 2M cells mL^−1^ and incubated at room temperature for 15 minutes. Cells were then centrifuged at 350g for 5 min and washed once with their respective culturing medium followed by resuspension in RPMI◻1640 medium (Life Technologies) containing 5% human serum (One Lambda, Los Angeles, CA, USA), 100 U mL^−1^ penicillin (Life Technologies), 100 μg mL^−1^ streptomycin (Life Technologies), and 2 mM GlutaMAX◻1 (Life Technologies), supplemented with 5 ng mL^−1^ IL◻7. The cells (100, 000 T cells, 50,000 mDCs or 50,000 T cells and 25,000 mDCs) were added to the middle well in the agar set up. In one of the outside agar wells 100 ng CCL19 was added to generate a gradient by diffusion (Figure 1A). In experiments where the diffusion was visualized, 100 ng Dextran, Texas Red, 10,000 MW, Neutral (Life Technologies) was added at the same time.

**Figure 1.**
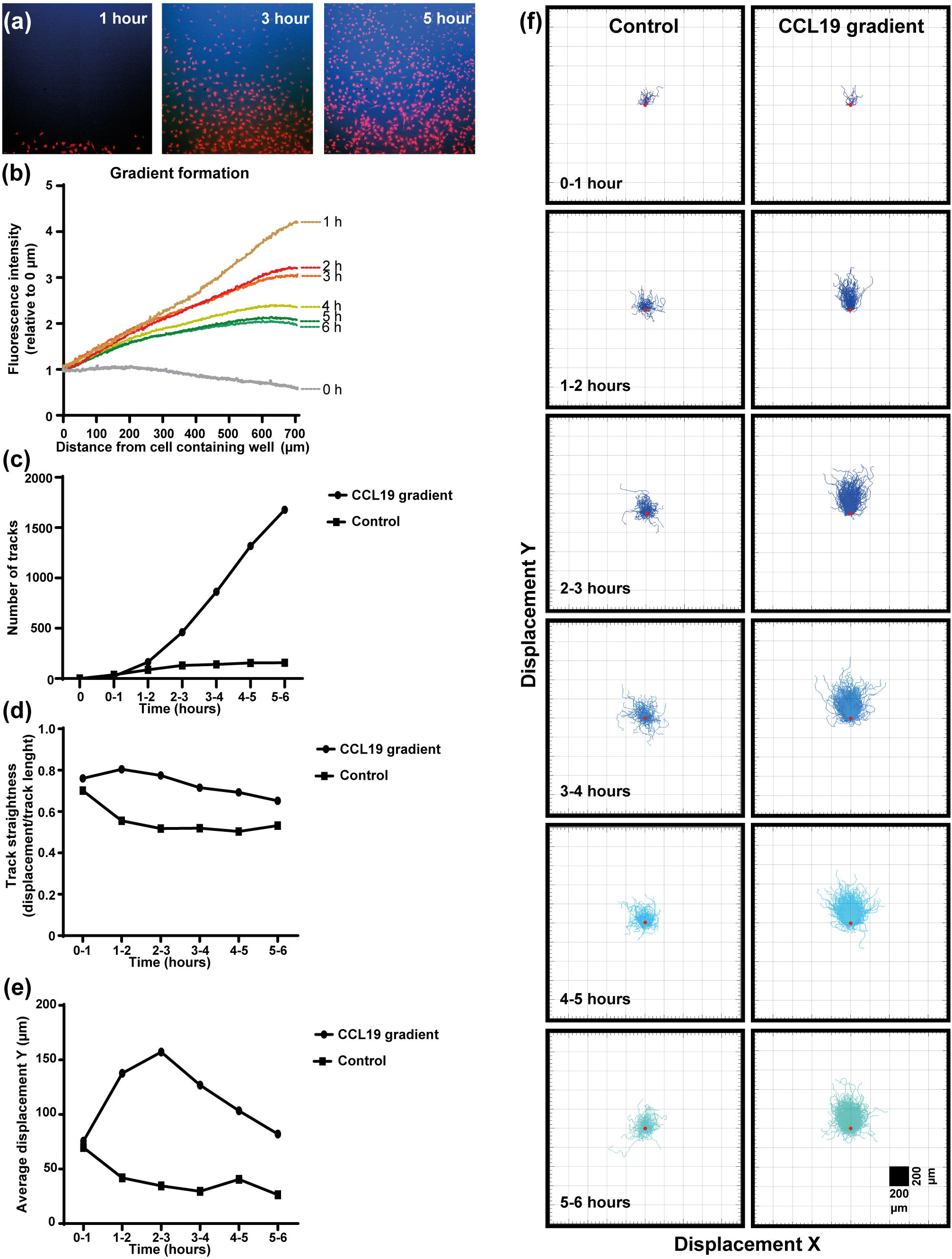
Dendritic cells respond chemotactically to a cytokine gradient in the “under-agar” assay, which lasts for several hours as shown by fluorescent dextrans. **a)** Images showing the diffusion of fluorescent dextran (blue) and migration of mDCs (red) at one, three and five hours of incubation. **b)** The formation of a gradient in the under-agar assay as shown by diffusion of 10 k.Da dextrans. **c)** Graph showing the number of tracked mDCs over time in response to a CCL19 gradient or no chemokine (control). d) Graph showing the track straightness ofmDCs over time in response to a CCL19 gradient or no chemokine (control). **e)** Graph showing the displacement of mDCs in the direction of the CCL19 gradient or in the same direction without chemokine (control). **f)** “Spider plots” showing the tracks ofmDCs over time in response to a CCL 19 gradient or no chemokine (control) plotted from a single origin point (red dot). All data is from one representative experiment that was repeated at least three times.

The μslide containing the agar and cells was then placed on an inverted Nikon TI-e (Nikon, Tokyo, Japan) and visualized using a 10× 0.4 NA Nikon lens and an Andor Zyla 5.5 camera (Oxford Instruments, Abingdon, UK). An image was taken every minute for up to the indicated times.

### Image analysis with Imaris

Imaris software (Oxford Instruments) was used to analyze the image sequences. Using the spot tracking module the cells were detected by their respective fluorescence label and tracked in 1-hour blocks.

### Statistics

Prism 8.1.2 (GraphPad, San Diego, CA, USA) was used for all statistical analyses.

## 3 Results

### Set up and gradient generation in an under-agar assay

In the under-agar set up used in this study (Supplementary Figure 1) fluorescent dextrans of similar size to CCL19 (~10 kDa) were used to measure the “steepness” and duration of the gradient in real-time concurrent with the mDC response to CCL19 (Figure 1, supplementary video 1). Measuring the diffusion of the dextrans indicates that a steep gradient is formed after one hour and that the gradient persists for up to six hours (Figure 1b). By adding CCL19 at the same time as the fluorescent dextrans it was possible to analyze the response of mDCs to the visual gradient that was formed. At one-hour post addition of CCL19 and dextrans, a “wave” of mDCs migrating out of the well and going under the agar can be observed. This matches the visualized diffusion of the fluorescent dextrans, which reached the mDCs in the middle well in the agar at that time. The increased directional migration of the mDCs compared to the control condition lasted for up to six hours, gradually getting less directional as the steepness of the gradient decreases over time, again matching the gradient as visualized by the dextrans. Analysis of cell movement highlights the difference in migration between cells that receive a CCL19 signal and those that do not. This allows the comparison of “classic” chemotaxis parameters such as the number of responding cells (Figure 1c), track straightness (Figure 1d), migration speed (Figure 1e), and the displacement in the X and Y direction of the cells (Figure 1f).

### Chemotaxis and chemokinesis can be clearly distinguished using time-lapse imaging

When our under-agar assay was used as an endpoint assay, in the same way as the original under-agarose assay published in 1975 (Nelson, Quie, and Simmons 1975) we can see that with both a gradient of CCL19 and a uniform concentration, migration of mDCs and T cells can be observed (Figure 2). Even though the cell number appears to be visually higher when there is a gradient present for the mDCs, chemokinesis can easily be mistaken for chemotaxis in this type of analysis. We also noted that visually there seemed to be increased migration by T cells when co-incubated with mDCs, suggesting some interaction between the cells or the tracks they make in the agar.

**Figure 2.**
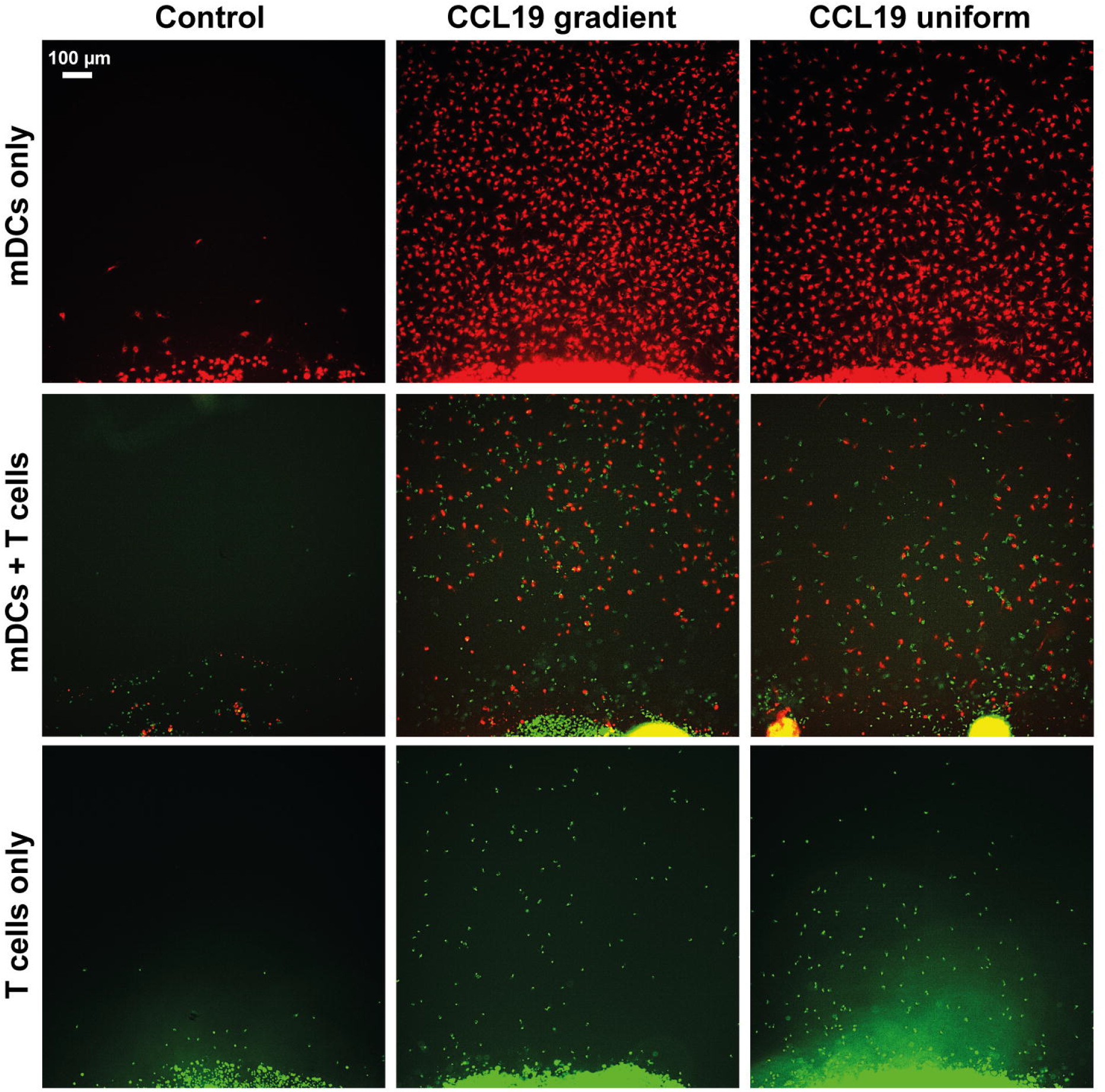
Static assays do not clearly distinguish between chemotaxis and chemokinesis. Images showing migration after 16 hours of incubation of mDCs (red) and T cells (green) without chemokine (Control), in response to a CCL19 gradient, and 100 ng mL-1 uniform CCL19 concentration.

When comparing a gradient of CCL19 with a uniform concentration of CCL19 in real-time, we observe apparent differences in the way mDCs are migrating (Supplementary video 1 and 2). When the tracks are plotted from a single origin point, we can see that mDCs display an evident chemotactic migration toward the CCL19 gradient compared to mDCs that have been exposed to a uniform CCL19 concentration (Figure 3). This mDC migration was unaffected by the presence of T cells during co-migration experiments (Figure 4). However, unlike mDCs, T cells do not show true chemotactic migration toward CCL19, either when migrating in the presence or absence of mDCs (Figure 4, supplementary video 3, 4, 5, 6).

**Figure 3.**
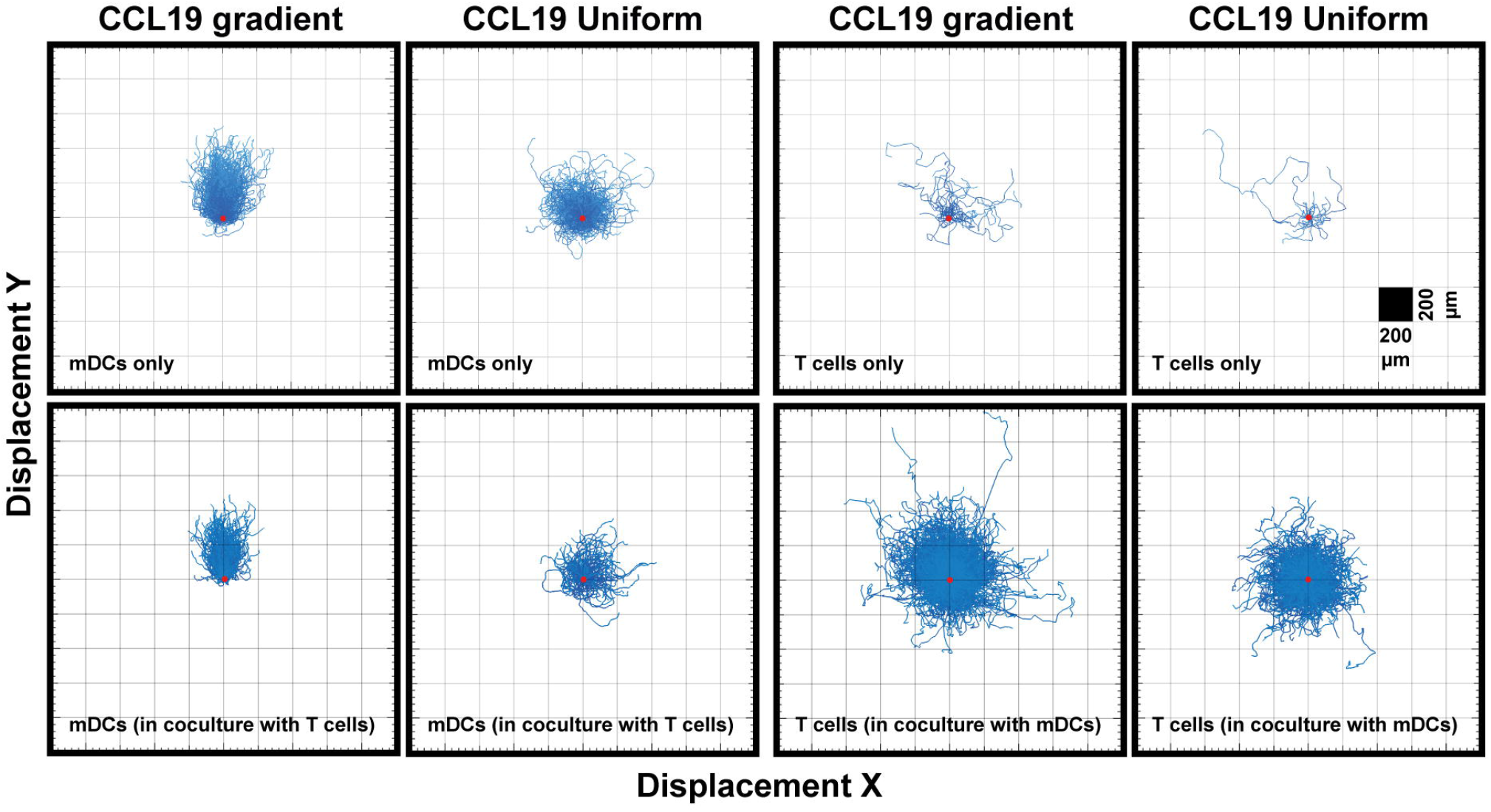
Unlike DCs, T cells do not show a strong chemotactic response to a CCL19 gradient. “Spider plots” showing the tracks of mDCs and T cells over time in response to a CCL19 gradient or a uniform 100 ng mL-1 CCL19 concentration plotted from a single origin point (red dot). A11 data is from one representative experiment that was repeated at least three times.

**Figure 4.**
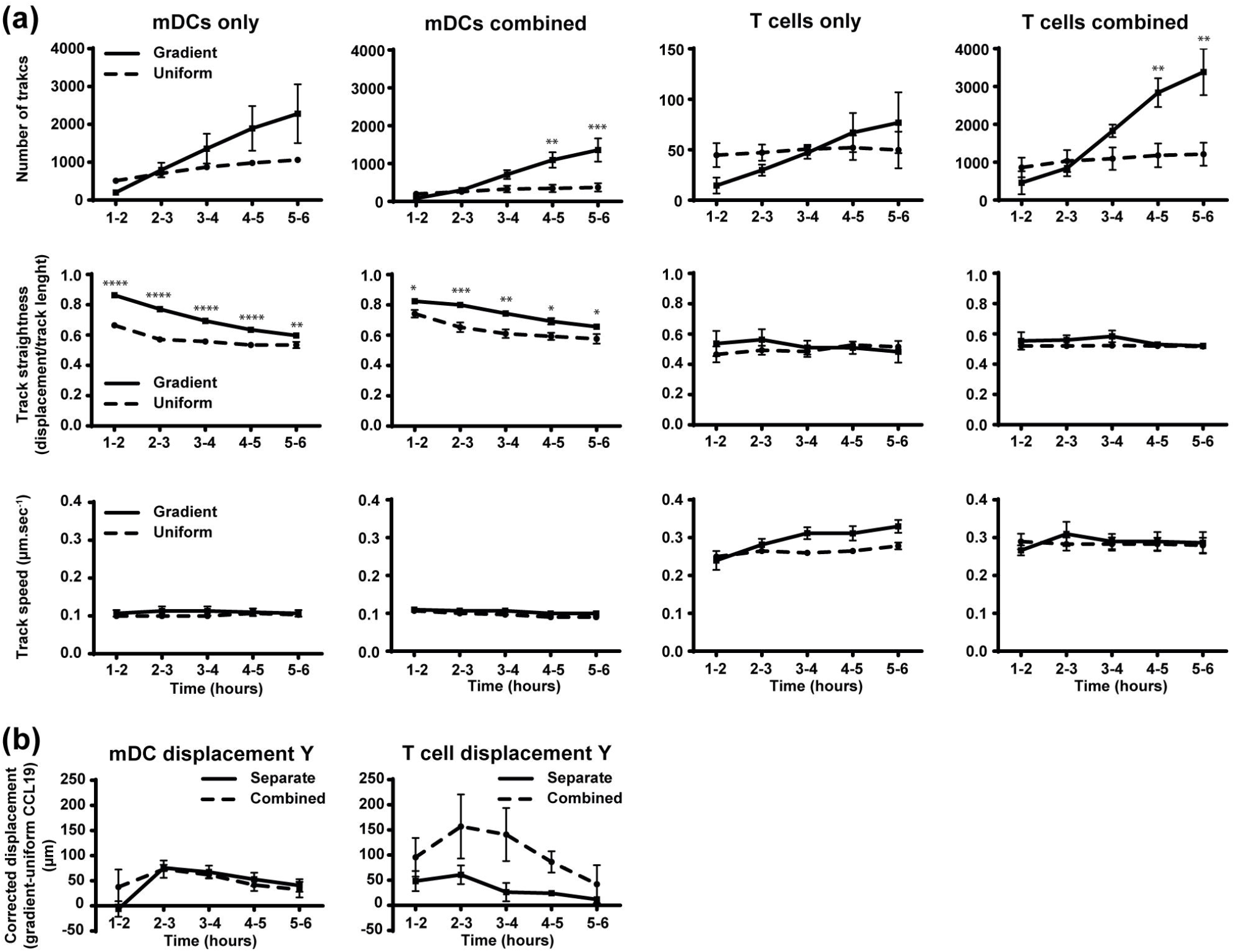
Migration behavior of DCs is not altered by the presence of T cells while the number of T cells and their displacement toward CCL19 is increased in the presence of mDCs. **a)** Graphs showing the number of tracked cells, track straightness and track speed of mDCs and T cells over time in response to a CCL19 gradient or uniform 100 ng mL-1 CCL19 concentration. **b)** Graphs showing the displacement ofmDCs and T cells towards a gradient of CCL19. The displacement was analyzed by taking the average displacement of T cells toward a CCL19 gradient and subtracting the average displacement of T cells in the same direction in a uniform CCL19 concentration_ Data are combined from three independent experiments and are presented as mean +/− SEM_Results were analyzed using multiple t-tests with the assumption that all populations have the same scatter and was corrected for multiple comparisons using the Holm-Sidak method. * P-value < 0_05, ** P-value < 0_01, *** P-value < 0.001, **** P-value < 0_0001

When the number of tracks, track straightness, and track speed were plotted and compared over time, a clear difference can be observed when comparing mDCs responding to a CCL19 gradient and a uniform CCL19 concentration (Figure 4a). Although there was an increase in the number of mDCs migrating in the absence of T cells with a CCL19 gradient, this was not statistically significant. The mDCs that are migrating do so in significantly straighter tracks when responding to the gradient. The track speed is similar between the two conditions, indicating that a gradient is less critical for inducing migration but that it regulates the direction of the mDCs. T cells, on the other hand, show only minor differences in migration to a gradient or a uniform CCL19 concentration (Figure 4a). However, when the displacement was analyzed by subtracting the displacement of T cells in a uniform CCL19 concentration from the displacement of T cells in a CCL19 gradient, there was an increase in displacement toward CCL19 (Figure 4b, solid line).

### Co-culture of DCs and T cells changes T cell behavior but not DC behavior

When T cells were cultured separately, very few cells migrated compared to the mDCs, suggesting some relative impairment of T cell migration. However, T cells cultured under the same conditions co-incubated with mDCs migrated ~30 times more frequently (Figure 4a), which refutes that notion. There is also an indication that there was an increase in displacement toward CCL19 (Figure 5b, dotted line), although, because of donor variation, this did not reach statistical significance.

## 4 Discussion

The data presented in this study showed that human polyclonal T cells, unlike mDCs, respond to a CCL19 gradient largely chemokinetic, showing a random walk with directional bias and showing similarities to the response to a uniform CCL19 concentration. Previously published data showed that the human Jurkat T cell line, transfected with CCR7 to respond to CCL19, responded chemotacticly to a 100 nM CCL19 gradient under experimental flow conditions (Wu et al. 2015). However, the authors of this study used a fibronectin-coated microfluidic device, and it has been reported that T lymphocytes can orientate their migration based on the direction of fluid flow during integrin-mediated migration (Valignat et al. 2013). Similarly, another study reported that T cells orientate towards 100 nM CCL19 gradient (Nandagopal, Wu, and Lin 2011), which is similar to what we see with the biased random walk. In our hands, T cells in a CCL19 gradient showed an increase in displacement toward CCL19. This suggests that CCL19 induces a bias toward a gradient of CCL19, similar to a biased random walk, in T cells. It has been previously suggested that T cells can display a super diffuse random walk (Lévy walk) that emerges from an explorative process, informed movement, and interaction with the environment (Krummel, Bartumeus, and Gérard 2016). These data are consistent with the behavior of T cells that use Lévy strategies to allow for optimal scanning of their environment for antigen (Harris et al. 2012).

When T cells were coincubated with mDCs there was a large increase in migrating T cells. The explanation is likely to be in the interactions of T cells with DCs. As visualized in the videos, T cells form brief contacts with DCs. This is consistent with the literature, showing that T cells briefly adhere to DCs, even in the absence of cognate antigen (Miller et al. 2004). This means that when a large population of DCs is introduced and is migrating directionally, T cell contacts with those cells will impart “momentum” in the same direction. The presence of mDCs did not alter the ‘random walk’ of the T cells. In contrast, mDCs did not show any difference in their migration behavior, whether tested in isolation or when co-incubated with T cells (Figure 5).

Our results suggest that T cells are programmed to respond differently to a gradient of CCL19 compared to DCs. This also indicates that T cells that are co-migrating with mDCs are not following pre-formed tracks made by the mDCs. This observation is further supported by the live-cell videos (supplementary video 5 and 6) where T cells can be seen migrating ahead of the mDCs. It has been reported that mature DCs can produce CCL19 that induces migration and scanning in T cells (Kaiser et al. 2005). This could be the reason for the increased T cell migration observed in our assays. However, further study is necessary to confirm this or if another mechanism is responsible for the increased T cell motility.

The random walk with a chemotactic bias that the T cells show toward a gradient of CCL19 observed using our live-cell assay fits with models that have been previously generated (Bogle and Dunbar 2012). Computational modeling of the random walk has shown that this movement pattern would be more efficient in activating T cells than chemotaxis alone (Riggs et al. 2008) as this enables efficient scanning of antigen-presenting cells in lymphoid tissues.

It seems likely that T cells’ ultimate paths will be determined by their inherent explorative behavior combined with their interactions with cells and molecules in their environment, given that they continuously scan and form transient cell-cell connections with professional antigen-presenting cells (Dustin 2008), that themselves can produce cytokines that stimulate T cell migration (Castellino et al. 2006; Penna et al. 2002; Beaty, Rose, and Sung 2007; Kaiser et al. 2005).

Using the assay presented here, it was possible to show a clear difference in migration behavior between mDCs and T cells. The T cell-specific motility patterns we observed highlight the exploratory behavior of T cells under CCR7 signaling that is likely to be crucial in generating optimal immune responses.

## Supporting information

Supplementary Video descriptions

## 6 Author Contributions

NPB and PRD obtained funding for the research. EJL, NPB and PRD designed the experiments. EJL carried out and analyzed all the experiments. HS obtained ethics for all work with primary human blood cells. EJL, HS and PRD wrote the manuscript.

