## Supplementary figures and images for "Live-cell microscopy reveals that human T cells primarily respond chemokinetically within a CCL19 gradient that induces chemotaxis in dendritic cells"

### Supplementary Figure 1

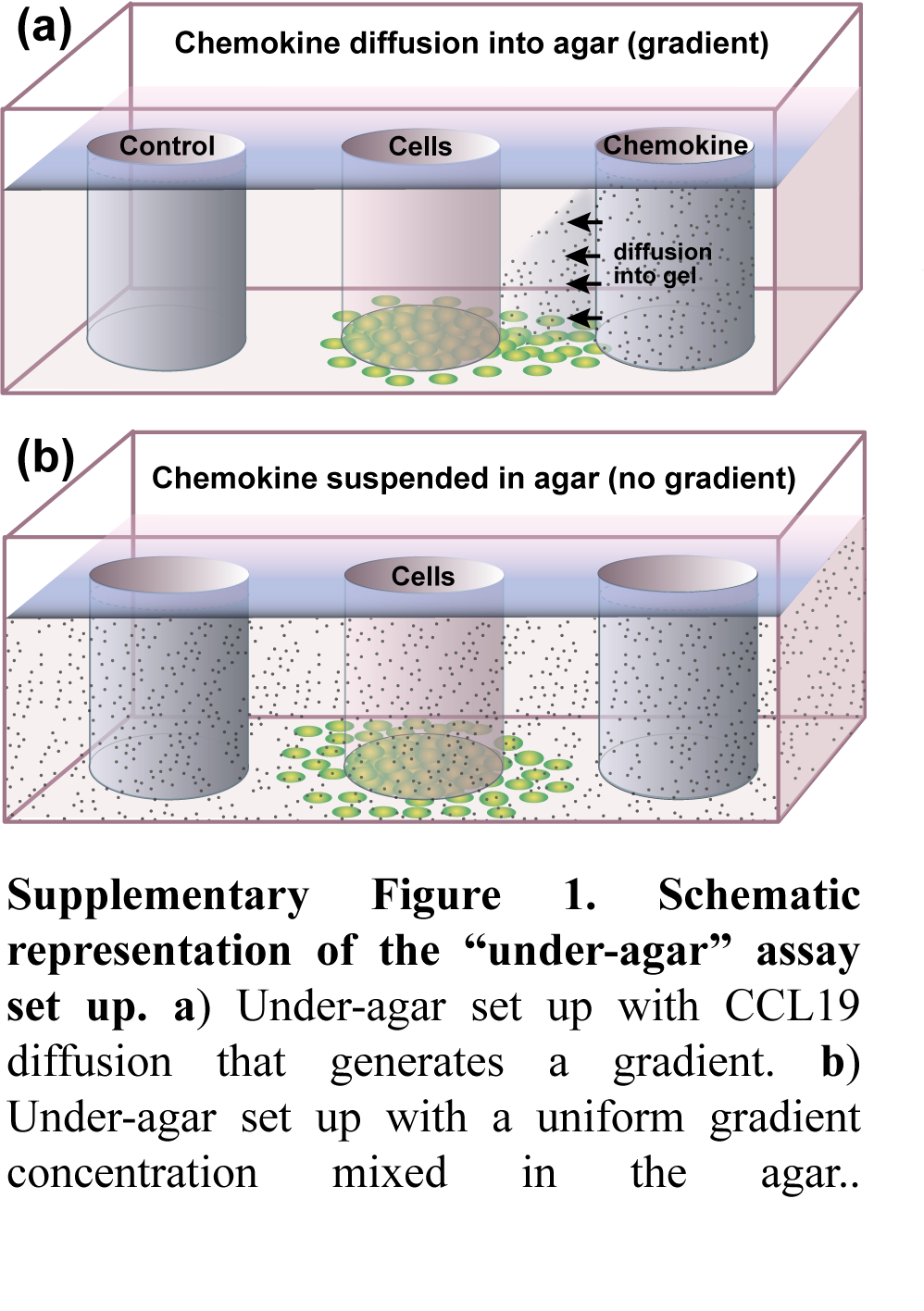
