## Supplementary Video descriptions for "Live-cell microscopy reveals that human T cells primarily respond chemokinetically within a CCL19 gradient that induces chemotaxis in dendritic cells"

Supplementary Material

**Supplementary video 1. Gradient formation and mDCs response to CCL19 in the under-agar assay.** 10 kDa fluorescent dextrans (yellow) were added at the same time as 100 ng CCL19 to analyze the diffusion and the migration of mDCs (cyan) in the under-agar assay in real-time.

**Supplementary video 2. mDCs migrating in a uniform CCL19 concentration.** Time-lapse showing the migration response of mDCs up to 6 hours after being added the under-agar assay containing a uniform 100 ng mL^-1^ CCL19 concentration.

**Supplementary video 3. T cells migrating to a CCL19 gradient.** Time-lapse showing the migration response of T cells up to 6 hours after 100 ng CCL19 was added to the under-agar assay.

**Supplementary video 4. T cells migrating in a uniform CCL19 concentration.** Time-lapse showing the migration response of mDCs up to 6 hours after being added the under-agar assay containing a uniform 100 ng mL^-1^ CCL19 concentration.

**Supplementary video 5. mDCs and T cells migrating in a co-culture to a CCL19 gradient.** Time-lapse showing the migration response of mDCs (red) and T cells (green) in co-culture up to 6 hours after 100 ng CCL19 was added to the under-agar assay.

**Supplementary video 6. mDCs and T cells migrating in a co-culture to a uniform CCL19 concentration.** Time-lapse showing the migration response of mDCs (red) and T cells (green) in co-culture up to 6 hours after being added the under-agar assay containing a uniform 100 ng mL^-1^ CCL19 concentration.
